# Improving on hash-based probabilistic sequence classification using multiple spaced seeds and multi-index Bloom filters

**DOI:** 10.1101/434795

**Authors:** Justin Chu, Hamid Mohamadi, Emre Erhan, Jeffery Tse, Readman Chiu, Sarah Yeo, Inanc Birol

## Abstract

Alignment-free classification of sequences against collections of sequences has enabled high-throughput processing of sequencing data in many bioinformatics analysis pipelines. Originally hash-table based, much work has been done to improve and reduce the memory requirement of indexing of *k*-mer sequences with probabilistic indexing strategies. These efforts have led to lower memory highly efficient indexes, but often lack sensitivity in the face of sequencing errors or polymorphism because they are *k*-mer based. To address this, we designed a new memory efficient data structure that can tolerate mismatches using multiple spaced seeds, called a multi-index Bloom Filter. Implemented as part of BioBloom Tools, we demonstrate our algorithm in two applications, read binning for targeted assembly and taxonomic read assignment. Our tool shows a higher sensitivity and specificity for read-binning than BWA MEM at an order of magnitude less time. For taxonomic classification, we show higher sensitivity than CLARK-S at an order of magnitude less time while using half the memory.

## INTRODUCTION

In computational biology, sequence classification is a common task with many applications such as contamination screening (1), pathogen detection (2), metagenomics (3), and targeted assembly from shotgun sequence data (4,5). Though this problem is addressable via sequence alignment (6), the scale of modern datasets (in both the scale of our query and reference sequences), has spurred the development of faster alignment-free hashed based similarity methods (3) as exact genomic coordinates are often unnecessary and leading to more computation than necessary. We have developed a novel probabilistic data structure based on Bloom filters (7) that implicitly stores hashed data (to reduce memory usage) yet can better handle sequence polymorphisms and errors with multiple spaced seeds, increasing the sensitivity of hashed-based sequence classification.

## Background

Using any hash-based methods for indexing sequencing data involves creating an incomplete representation of the data. The most common alignment-free indexing methods are *k*-mer (substring of length *k*) based. These methods work by breaking a reference sequence into *k*-mers and indexing them (often in a hash table). To classify sequences, they are also broken into *k*-mers and queried against the index to check for shared *k*-mers. If a significant number of *k*-mers are found within the reference set, the read is then classified. The *k*-mers must be long enough such that they are unlikely to be the same between indexed targets, especially if there is substantial sequences similarity.

However, *k*-mers cannot compensate for differences between references and queries that occur within *k* base pairs of each other. This limitation of *k*-mers has motivated us to use spaced seeds (1) (also called gapped q-grams (2)).

Spaced seeds are the current state-of-the-art for approximate sequence matching in bioinformatics. They are a modification to the standard *k*-mer where some positions on the *k*-mer are set to be “don’t care” or wildcard to catch the spaced matches between sequences. They were originally proposed in PatternHunter in 2002 (1) and have been increasingly used since then to improve the sensitivity and specificity of homology search algorithms (3–7). Employing *multiple* spaced seeds together can greatly increase the sensitivity of homology searching (8). Spaced seeds have been employed in metagenomics studies and successfully performed to improve the sensitivity of classification in metagenomic classification (9,10).

### Probabilistic data structures

Probabilistic data structures (11) are a class of data structures that focus on representing data approximately, so query operations can sometimes produce an incorrect result. The use of probabilistic data structures in bioinformatics has expanded in recent years, owing to their speed (constant hash table-like speed) and low memory usage. However, the use of these data structures is a double-edged sword due to the existence of false positives.

Depending on the application, the permissive rate of false positives is constantly re-evaluated with a multitude of different mitigation strategies proposed. No matter the methods, at the core of false positive reduction strategies is the use of reduction of base probabilities and the use of conditional probability from independent events. For example, from the first probabilistic data structure, the humble Bloom filter (12), false positives are reduced by lowering the occupancy of the Bloom filter (decreasing the base probability of a false positive) or by increasing the number of hash function used (exploiting the conditional probability of multiple independent events). These principles have not changed; however, it is important to note that there may be aspects of the data type being indexed and variations of the data structure being used that are underutilized when attempting to reduce the false positive rate. For instance, methods that utilize probabilistic data structures for key-value associations in sequence analysis often consider every inserted key as an independent event (13,14). Conceptually, this manifests itself in assigning a false positive rate (FPR) for each key query performed. However, in the biological sciences, unlike in many applications of these data structures in the computational sciences, each key is actually some kind of decomposition of parts of the same sequences (*e.g.* a *k*-mer) and thus not independent. This pseudo-redundancy can be exploited in to reduce the relative FPR when querying the data structure (15).

The use of Bloom filters for sequences based classification was originally developed in the tool Fast and Accurate Classification of Sequences (16). We later developed BioBloom Tools (BBT) (17), which used heuristics to optimize speed and reduce the effect of false positives. Though BBT proved effective when using a small number of filters, determining the specific reference of a queried sequence requires multiple Bloom filters, which can lead to an O(*n*) time complexity where *n* is the number of references. Here, we have extended the functionality of Bloom filters for the classification against multiple references (key-value association) in a novel data structure called a multi-index Bloom Filter (miBF). While miBF shares similarities with existing data structures, it has properties that allow it to synergize with spaced seeds.

### Related data structures and algorithms

Bloomier filters are first probabilistic data structures that allowed key-value associations, utilizing perfect hashing to prevent collisions between values (18), though has seen no prominent use in bioinformatics; Marchet *et al* suspect that this may be that no free implementation exists for this data structure as of yet (14). At any rate, Bloomier filters have been superseded for applications in bioinformatics by higher performance data structures, such as the quasi-dictionary (14,19) and Othello data structure (13,20).

The quasi-dictionary allows for key-value lookups and was originally proposed for applications in metagenomics(19). The quasi-dictionary uses bbHash (21), a minimal perfect hashing scheme that only requires 3 bits per element. Though this solves issues for preventing collisions of values, false positives are still an issue. To reduce false positives, a small value (in practice 12 bits, derived from the key is also stored in the indexed position, a strategy likely derived from quotient filters (22) where the key-derived value is referred to as a “quotient”. Assuming 16-values, each values will take 12+3+16 = 31bits per element. Similar data structures to this include a compact coloured de Bruijn graph implemented in Pufferfish (23) (which also uses bbHash) and works on some very similar principles.

The Othello data structure was originally designed for network forwarding lookup optimization (20) but has now seen uses in metagenomics (13) and even RNA seq analysis (24). It bears some resemblance to another probabilistic data structure called Cuckoo Filter (25). Like Cuckoo filters, it works by using 2 hash functions for 2 potential hash locations, however, instead of storing a quotient of some sort in 2 possible locations, the value is stored in 2 tables, where the value stored is the exclusive-or of 2 hash values. Just like Cuckoo filters, the second hash value used is a way of dealing with collisions. When a collision occurs that is unresolvable (both hash locations are use) the element cannot be inserted but in practice only results in a few missing elements. The memory usage is O(*ln*), where *l* is the size of the value type stored and *n* is the number of elements stored. They require that each table be at least 1.33*n* in size, thus assuming 16-bit values, each value will cost (16+16)×1.33 = 43 bits per element. They refer to false positives as alien *k*-mers but use a sliding window approach where adjacent *k*-mers are evaluated, similar to the method presented here to minimize false positives.

Our miBF belong in the same family of these data structures, with similar theoretical performance and properties, but is slightly less suitable for *k*-mers. This is due to our strategy of handling hash collisions and dealing with false positives. At suggested parameterization, our miBF requires the least amount of memory per key compared to these methods (less than 20bits, assuming 16bit values and that every single spaced seed in a set of multiple spaced seeds is considered its own key), however such a statement is somewhat meaningless without an expected FPR. Because the FPR for a single key query of miBF is contingent on the value pair it classifies to, a directed comparison to other methods is difficult. In short, miBFs are not designed to be used for single queries. To most effectively use miBFs, they must be used with multiple queries over the same sequence, effectively amortizing the FPR to a small value. Because of this, we consider the FPR to be a function of not only the miBF itself but the length of the sequence being queried.

## MATERIAL AND METHODS

### Multi-Index Bloom filters vs Bloom filters

We developed a novel Bloom filter-based data structure that can perform constant time key-value associations called multi-index Bloom Filter (miBF). For standard Bloom filters, elements of a sequence, such as *k*-mers, are queried to determine whether they are or not members of those decomposed from the reference set. Like Bloom filters, the memory usage of miBF does not depend on the size of the *k*-mer or the spaced seeds used. With a Bloom filter, querying for the set of origin between multiple reference sets requires the construction and use of multiple Bloom filters leading to O(*n*) time complexity when querying, where *n* is the numbers of reference sets/filters. Unlike Bloom filters, querying for the set of origin for an element requires only one data structure and is performed in constant time.

### implementation details

The miBF data structure is implemented as part of BioBloom Tools (BBT) (https://github.com/bcgsc/biobloom), implemented in C++ with components and our BTL Bloom Filter (https://github.com/bcgsc/btl_bloomfilter). For space seed hashing we use a modified version of ntHash (26) a recursive rolling hash specialized for nucleotide sequences. Finally, we use components and algorithms from the C++ Boost libraries (27) and the Succinct Data Structures Library (28).

### Spaced seeds

To better utilize the available memory relative to false positive rate, Bloom filters can use multiple hash functions. However, there is no requirement for the hash values to be derived from the same *k*-mer. We have adapted our miBF to hash multiple spaced seeds instead of traditional *k*-mers. Naively, one may simply insert each spaced seed as its own element (using multiple hash functions for each insertion), yet, since the seeds in the same “frame” of the sequence are dependent we can instead use a set of spaced seeds in the place of multiple hash functions. By allowing for some spaced seeds in a frame miss, we can better tolerate mismatches when classifying sequences. By default, we will accept all but one seed in a frame to miss, but it is possible to change this and to help decrease the FPR, at the cost of sensitivity. There is no restriction on length or weight (number of required match positions) for spaced seeds used in our data structure, however, to enable us to store only one compliment of the sequence, we use spaced seeds that either share mirrored match positions to another seed or is palindromic. This allows us to save memory by storing each sequence only once and not both forward and reverse-complement, analogous to only storing one canonical *k*-mers (comparing both the forward and reverse-complement of a *k*-mer and consistently picking one).

### miBF Structure

Conceptually, the miBF can be thought of as three separate arrays. One behaves like a traditional Bloom filter bit array, storing the presence or absence of an element in the set (Fig. 1A). The next stores the rank information of the bit array at specific intervals; this allows for constant time rank information access (29) to positions on the bit array. The third ID array stores integer identifiers for each element in the bit array. These integer IDs can be used to represent any arbitrary classification category.

**Figure 1.**
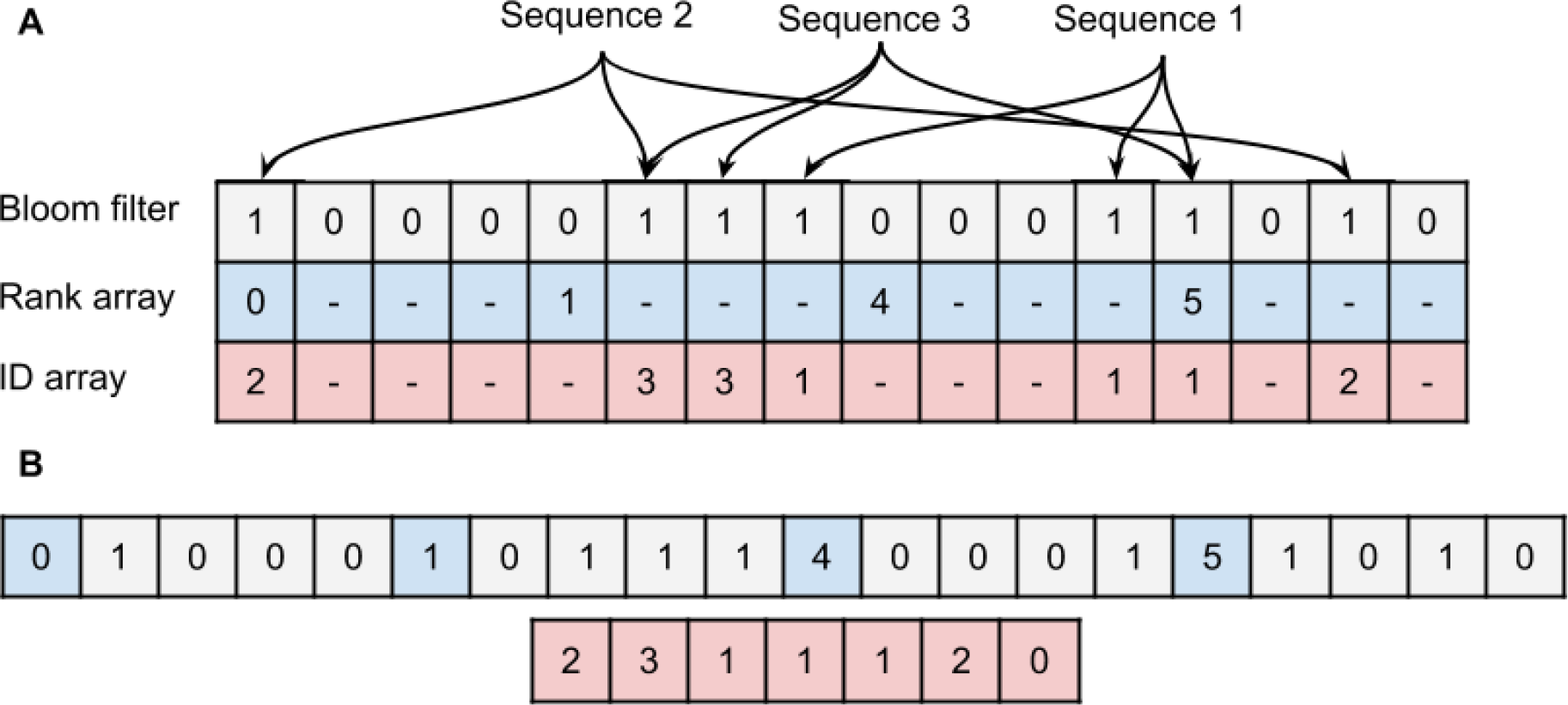
A multi-index Bloom Filter data structure visualization. A: Visualization of 3 tables used to represent the miBF and how they interact. B: Visualization of the true form of the data with an interleaved form of the bit vector. The interval for the rank array is much larger than shown here (4 vs 512 bits) which reduces its overhead to 64/512 = 0.125 bits per position. Under our scheme, hash collisions are permissible and the chosen ID to be stored minimizes the number of removed seed values per sequence frame.

To query a single value, we first look up the hash value in the Bloom filter, if it hits we use the rank array in conjunction with the Bloom filter to determine the rank of the value, and finally, we look up the corresponding integer identifier in the ID array. To improve cache performance the Bloom filter and rank arrays are interleaved into a single data vector (30).

### Data structure construction

Construction of the miBF consists of multiple passes through the sequence set being indexed. To minimize memory usage overhead, all data is streamed, but multiple threads can be used on different parts of the data (so long as they are part of different IDs) at a time to improve runtime. This pass through the data populates the Bloom Filter, with the rank array is created afterwards, and subsequent passes are used in populating the ID array. The number of passes needed to construct a miBF is 2 + *h* where *h* is the number of hash functions.

Due to shared sequence or hash collisions, inserted values into the ID array may collide causing a loss of key-value association information; however, there are ways to insert values in such a way that we minimize any loss of data across sets of sequences. For every colliding ID, we attempt to ensure that at least one of the hash values of a frame (i.e. same position of the sequence) will contain that ID. This is done by populating the ID array in steps (Figure 2). When populating the ID array, we pass through the sequences *h* times in a stepwise fashion. In the first pass, we only allow 1 value of a frame to be inserted, in the second pass 2 values, etc. This way we can ensure that we values are fairly distributed in a streaming fashion.

**Figure 2.**
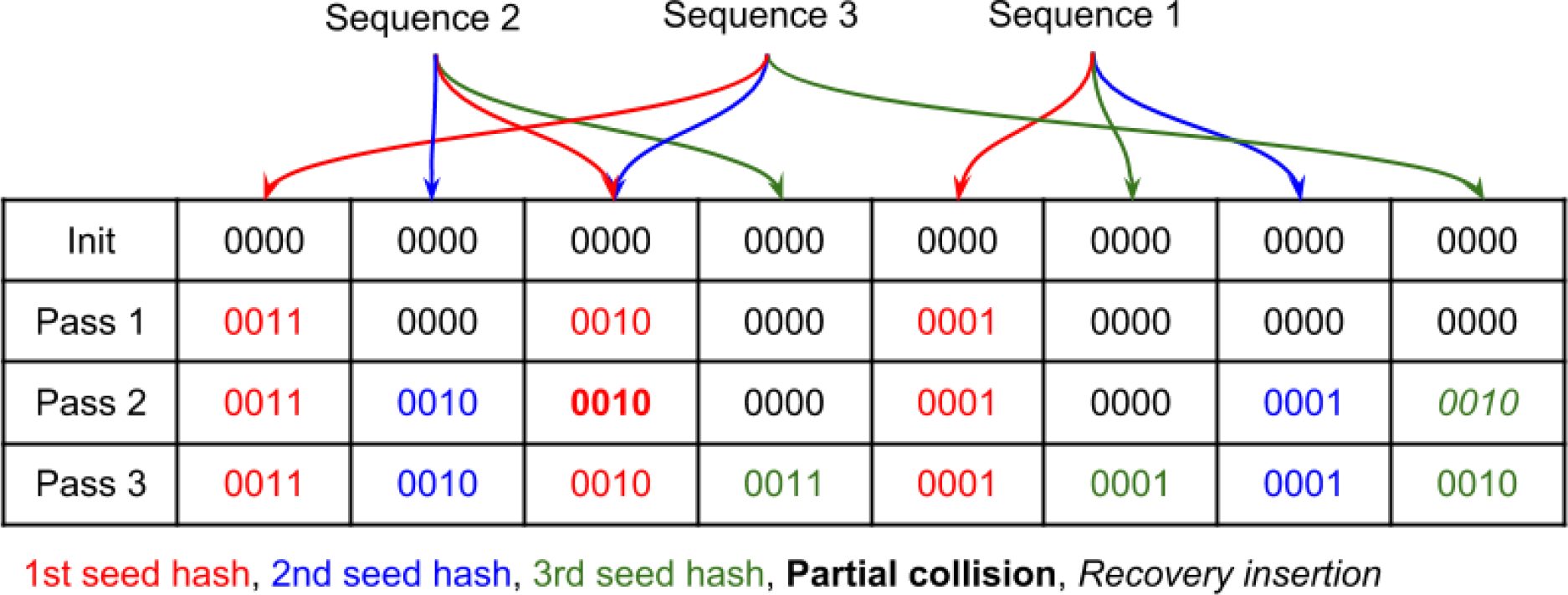
An example set of stepwise insertions into the ID array of the miBF. When partial collision this means a value could not be inserted in the first try within a step and had its next value placed in another location as a recovery insertion. We have highlighted the hash function value in different colours to help illustrate what value is being inserted at each step, but we note that there is no requirement for hash functions to be inserted in a pre-set order (random order is used in practice).

If the frame has values cannot be inserted (which can only occur in the first pass into the ID array) we set the value as saturated by reserving the first bit of the ID integer to denote saturation (Figure 3). The saturation bit lets us know if a key-value association information within a frame is lost, without destroying key-value pairs that are important for the classification of other sequences. After the first pass of insertions, we perform an additional pass to randomize the IDs in the saturated positions to minimize bias towards sequences inserted first. We do this by using a temporary array that stores occurrence counts per ID whilst streaming in conjunction with reservoir sampling (31). In this vector, we also store critical saturated positions needed for non-fully-saturated frames to ensure that we do not lose any information in these frames when randomizing the IDs. Finally, to further help minimize bias (Supp. Note S2) towards sequences inserted first, we will insert the next passes in the reverse direction, switching back to forward for the pass after that, until fully populated.

**Figure 3.**
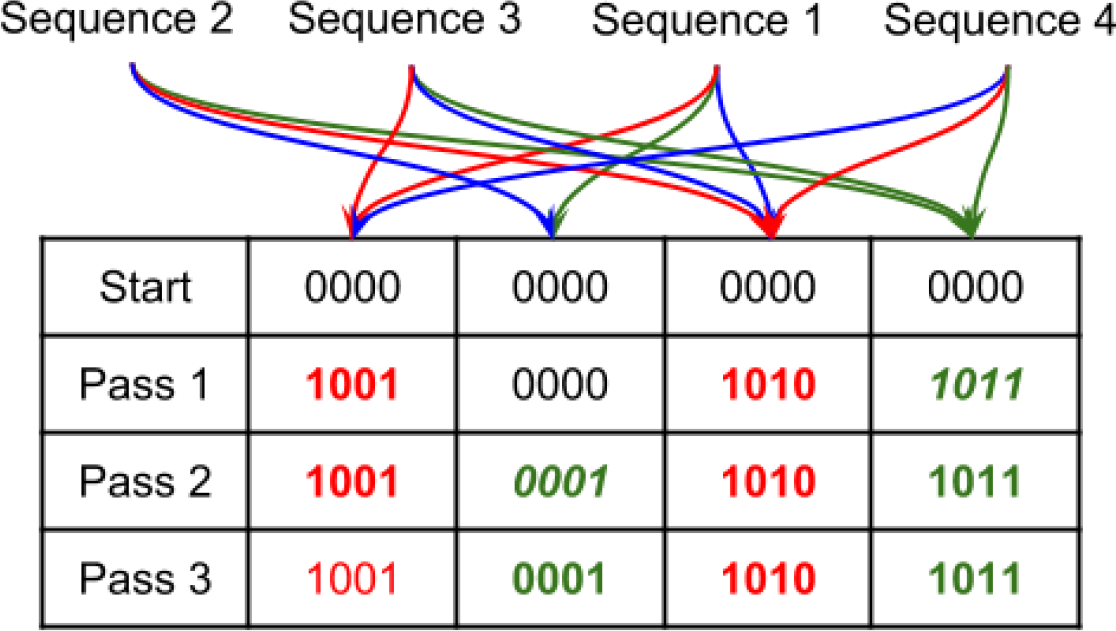
An example set of insertions causing saturation of some of the key-value association in the miBF. Sequences 1 and 2 can still be reliably be used for their key-value pairings, but sequences 3 and 4 will return completely saturated IDs such that they cannot be trusted to return all possible associated IDs to that sequence if it is queried.

### Queryin & Classification

A single lookup of the data structure is performed by hashing a set of spaced seeds in a frame and confirming if the value(s) are present and if present, using the rank value to retrieve the ID from the ID array. In this scheme, one can end up with multiple IDs for the query, with a high likelihood of one of them being a false positive. This may seem to invalidate the purpose of the data structure, however, one may observe that although false positive hits are independent, the hits to multiple frames of the same sequence will not be. Thus, to query reliably, we use adjacent frames to reinforce and reduce the effective FPR. This is similar to other methods is shown to help reduce the effective false positive rates in Bloom filters (15), with the difference of multiple indexes being considered.

Using adjacent sequences to minimize false positives can be generalized to calculate the FPR for any length greater than *k*. When classifying a sequence the FPR for each frame can be thought as a series of *n* independent Bernoulli trials (number of frames) so we can model the overall chance of a false positive using a binomial distribution. Conceptually, under this methodology, the data structure no longer has a fixed FPR; indeed, the chance of a false positive depends on the length of the sequence being queried, the length of the seed used, the FPR of the Bloom filter, and the frequency of each ID in the ID array.

We perform 2 stages in our classification. First, we determine if the matches to a particular ID are enough to determine the read is not a false positive match determined by a preset minimum FPR (1/1010 by default). Then we rank the significant candidate IDs and try to filter in additional candidates that match strongly enough that they can be considered a multi-match.

### Calculating the FPR of sequences of arbitrary length

As mentioned, we calculated the chance that a read is false positive by considering the entire sequence rather than a single lookup. The FPR of a Bloom filter is already well formulated (12), and if the occupancy of the Bloom filter, *b* is known it can be formulated as follows:

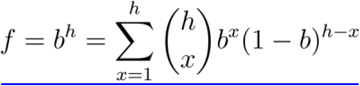

Where *h* is the number of multiple spaced seeds (traditionally number of hash functions) used for a single frame in the sequence. However, as we are also allowing some misses due to our use of multi-spaced seeds, the formulation becomes:

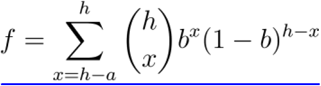

Where *a* is the number of allowed misses for the set of spaced seeds in a frame. As mentioned when classifying a sequence the FPR for each frame can be thought as a series of *n* independent Bernoulli trials so we can model the overall chance of a false discovery using a binomial distribution. That is, for a simple Bloom filter the chance of falsely classifying a sequence is easily determined by computing the cumulative density function (CDF) of the number of matches *m* - 1 and inverting it:

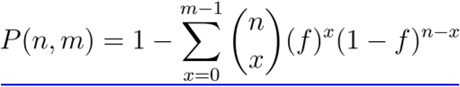

This will help in understanding how classification in a miBF is calculated for a single index. Computing the FPR of miBFs has similar formulation but incorporates the fact that you can use multiple indexes to help further reduce the FPR, though it should be noted that each index *i* is an additional test that you must consider and be corrected for. Before we compute the overall probability on an entire sequence for a given index *i*, we must first formulate the probability of falsely matching a frame of classification:

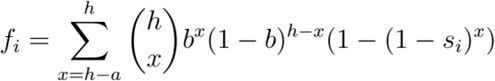

Where *s_i_* is the frequency of index *i* in the data array of the miBF. Thus, the overall probability for false classification for index *i* is:

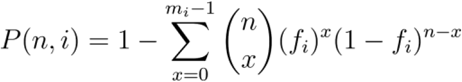

Out of all our tests for each index *i* we can take the best candidate (lowest p-value) and perform a multiple testing correction. In our implementation, we perform the Bonferroni correction (32,33):

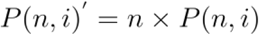

Because we expect a discrete pseudo-uniform distribution of p-values, and because common multiple testing correction methods (Supp. Fig. S2) are expecting a uniform distribution of p-values, these correction methods seem to overly conservative to the true false positive probability (which may lower the statistical power of the method). However, we believe this principled approach of calculating false positive rate on sequences can help improve the robustness and reproducibility of the method as these formulations ensure a minimum false positive rate to their results. Similar probabilistic classification methods in the past tend to use arbitrary scoring functions and thresholding, which can reduce the robustness of the method when sequence lengths or single element FPRs change.

### Determining and ranking multi matches

We filter out only reads that pass a minimum FPR threshold during classification. These reads are now considered not false positives, though may be associated with more than one candidate ID. We now need to determine if the classified reads are the best hit and if we can confidently say that the mapping is not shared between multiple hits ambiguously (due to shared sequence or close sequence homology). To rank these candidate hits, we use the following hierarchy:

1. Non-saturated frame counts: ID counts to frames that do not have any saturated IDs in the same frame, only counted once per frame
2. Non-saturated solid frame counts: ID counts to frames without saturation and only if the frame contains all seeds needed for a match, only counted once per frame
3. Frame counts: ID counts to frames regardless of saturation status, only counted once per frame
4. Non-saturated frame counts: ID counts that are not saturated, only counted once per frame allowing for frames to have some saturated IDs
5. Total non-saturated counts: all ID counts that do not have any saturated IDs
6. Total counts: all ID counts regardless of saturation status

Empirically, we found the non-saturated frame counts to be the best metric as it takes into account frames that may be repetitive (not including them into the predictions) yet is still able to tolerate mismatches (providing more evidence for classification). The frame counts are used to determine false positive hits (using the formulation in the previous section), and is generally a fairly reliable metric for ranking multi-matches but can be distorted if the sequence contains repetitive sequences. Total non-saturated counts are generally highly accurate but difficult to obtain a high number observation for and thus is less reliable in some case. The remaining metrics provide more observations to reinforce the confidence of a match, but because we assign ownership of an ID per frame and not spaced seed during construction, they are less useful at determining the best match of a sequence. All of these metrics should, in an ideal situation, agree with each other (always be higher if in the better match) and generally do in our tests. However, we also provide an option (−b) to filter classification of these cases where the metrics do not agree, improving the specificity, at the cost of a minor loss of sensitivity.

By default, we consider candidate ID matches to be too close to call if their non-saturated frame counts are within a threshold (−m = 2). The remaining candidates are returned, ranked by the hierarchy above, though it is likely that all remaining matches are likely homologous to the sequences in some capacity. Sources of error for all these counts come from repetitive sequences, homologous indexed references, sequencing error, polymorphism and random noise due to hash collisions during miBF construction.

### Speed optimization and heuristics

Because what is commonly classified is of a fixed length (i.e. read sequences), we can simply compute a fixed significant match threshold for each index upon seeing the first instance of a sequence of a particular length. By applying our Bonferroni-corrected critical p-value on this length for each index we compute the minimum number of frame counts needed to not be considered a false positive provided by the C++ Boost libraries (27) using the quantile function (34). With this, although optimal classification results are not guaranteed, the classification may now terminate early to improve throughput in practical situations. This situation occurs if the *si* values are sufficiently small and if the sequences are mostly independent. That is, if the contribution to reducing the FPR is dominated by *si*, we can achieve near-optimal classification when terminating classification early. This further increases the utility of the algorithm when one use utilizes many indexes rather than a few as this will cause *si* of each index to decrease accordingly. We terminate early through a parameter (−r = 10, by default allowing at least 10 frames of unambiguous significant matches until early termination) which we use in all our tests unless otherwise stated. The largest gains will be seen if there is low ambiguity (fewer candidate IDs, less sequence sharing) when classifying the sequences. This heuristic has no effect on the FPR, and only affects the accuracy of multi-match assignments.

## RESULTS

### Filtering reads for targeted assembly - Comparison with BWA MEM

The targeted assembly allows one to improve the throughput and reduce the complexity of assembly for applications where quick answers are helpful, such as clinical diagnostics for structural variants or other mutations. A typical procedure when performing targeted assembly is the extraction of sequencing reads in the target loci, before using these reads in a *de novo* assembly pipeline. This can be done via alignment or sequence classification since exact genomic coordinates are not necessary as the reads will be used in a *de novo* assembly afterwards. For this application, we compared the binning of reads with BWA MEM (v0.7.17) (35) with our method on a set of simulated reads. We simulated Illumina reads with depth of coverage ranging from 10x (229,800 read pairs) to 100x (2,303,019 read pairs) in increments of 10 using pIRS (v1.1.1) (36) from a gene set composed of 580 COSMIC (v77) genes (37) (targets) and an equal number of non-COSMIC genes randomly selected from RefSeq (38). We then indexed the set of 580 genes into a miBF using a set of 4 spaced seeds (Supp. Note 1).

These tests were performed on the same machine Intel Xeon E7-8867 2.5GHz with 64 threads. Compared to BWA-MEM, BBT obtained a higher overall sensitivity (99.9% vs 98.7%) and lower overall FPR (0.3% vs 0.4%) (Figure 4). We note that any multi-mapping reads are considered as part of multiple genes they map to, thus, if a sequence is shared, it will be considered a false positive for the shared but non-origin gene. We compared the runtime of each tool, finding that BBT runs 14x faster than BWA MEM on default settings with 64 threads. Memory usage of BBT on these set of 580 genes was 20MB.

**Figure 4.**
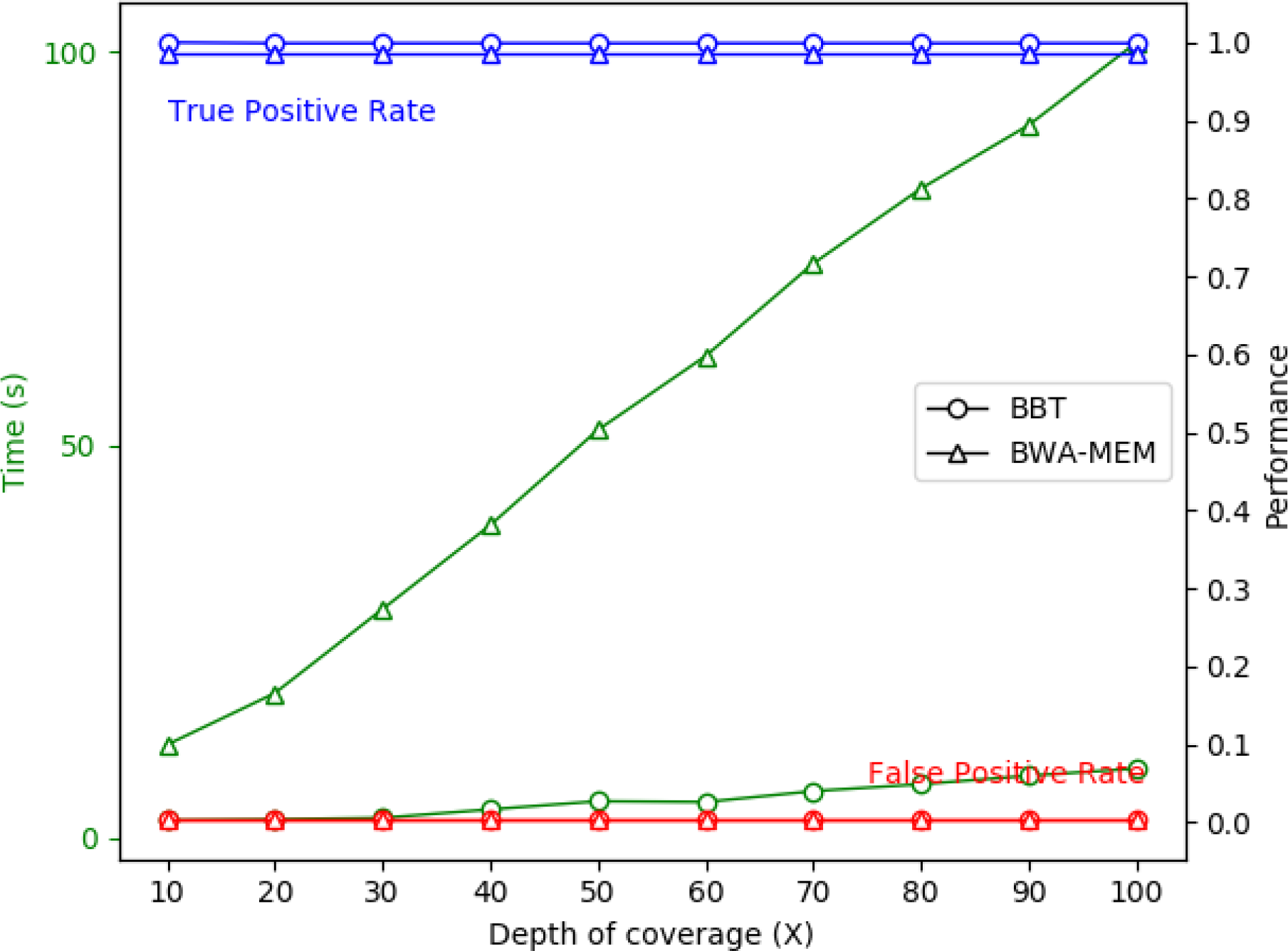
Read classification by BBT vs BWA-MEM alignment. 100bp simulated Illumina reads were simulated by pIRS (v1.1.1) with coverage depths ranging from 10X to 100X from 580 COSMIC (v77) genes and an equal number of non-COSMIC genes randomly selected from RefSeq. Read classification performance (right Y-axis) and run-time (green, left Y-axis) of BBT were compared against BWA-MEM The classified target of every read was compared against its true gene origin to calculate true (blue) and false positive rates (red).

**Figure 5.**
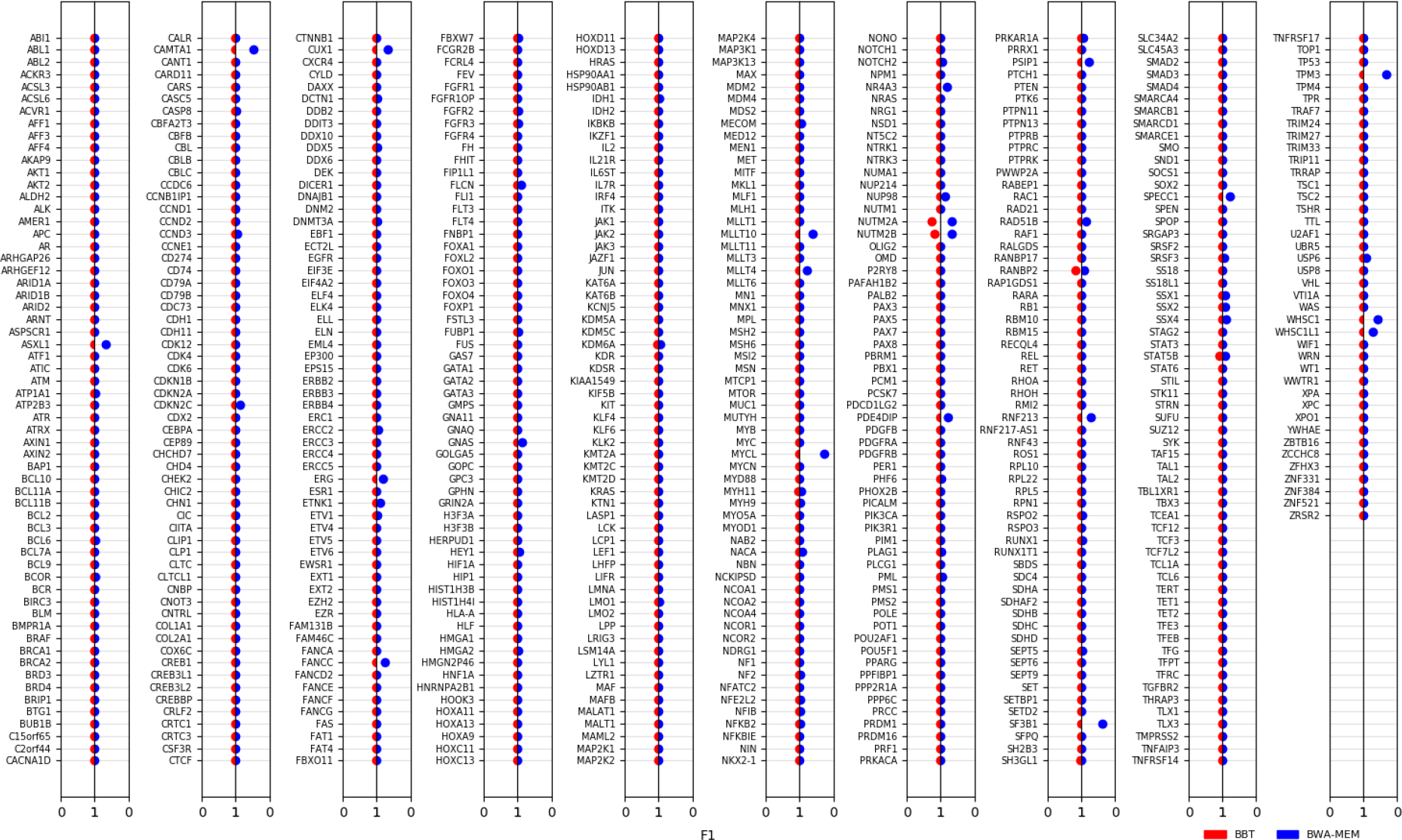
Per-gene comparison of classification performance by BBT vs BWA-MEM. F1 scores for both methods using the same simulation dataset described in Fig. 5 is calculated for each gene and plotted on the same horizontal line (red=BBT, blue=BWA-MEM). The scale on the X-axis for BWA-MEM on the right is reversed for easy visual comparison such that higher scores for both methods localize to the middle while lower scores are off-center.

On a per gene basis, BBT outperforms BWA-MEM in terms of F1 score, except for RANBP2, likely due to its similarity to RANBP17. This suggests that BWA-MEM in some cases may be better at discriminating between similar hits, whereas BBT will instead apply classification to both genes. Under default parameterizations, BBT is shown to be more sensitive at detecting homology than BWA-MEM but is in some cases less able to confidently assign them to a single target given closely related homologous sequences (more multi-mapping classification). This behavior is somewhat expected considering the use of minimal exact matches in FM-Index based alignment approaches compared to a fixed length spaced seed.

We also compared the memory and time usage of indexing but because the indexed set of COSMIC genes was very small it is difficult to compare the indexing methods. Instead, to compare the scalability of indexing we indexed a 3.5G fasta file consisting of ~1000 bacterial sequences. BWA took 1.5 hours to index in entire file while BBT took 9 hours to index the same file but only 1.5 hours using 16 threads. Normal BWA indexing cannot be multi-threaded.

### Metagenomic classification

Although BBT is a generic classification tool, when given the proper reference sequences it is possible for it to be used as the workhorse for a metagenomics classification pipeline. To demonstrate this, we compared BBT with CLARK (39) and CLARK-S (40) (the spaced seed variant of CLARK). We compared CLARK-S because it is the only metagenomic classification tool that we know of that supports multiple spaced seeds used to improve classification sensitivity. We also compare the method to CLARK because it is the predecessor to CLARK-S and well characterized against other tools as seen in other studies (41). Finally, like CLARK/CLARK-S, we can currently only construct databases for one taxonomic level at a time (unlike tools like Kraken (42)), thus facilitating more direct comparisons.

We first generated a standard database (bacterial + viral genomes, constructed May 2018) for both CLARK-S (using default seed set) and CLARK. To make sure the comparison performed were comparable, we used the same references sequences and taxon IDs that CLARK uses to construct our miBF (Supp. Note S1). To index the bacterial and viral genomes, CLARK took 24h, CLARK-S took 24.5h and BBT took 19 hours to generate an index. We requested 24 threads for each tool but CLARK generally did not use more than 1 CPU at a time, while BBT used around 12 CPUs on average, suggesting there is still room for optimization and better parallelism to be exploited when indexing for both tools.

There are some differences between our miBF index and CLARK/CLARK-S databases beyond our implicit representation of our seeds. First, we note that unlike CLARK-S and CLARK, seed sequences shared between different taxa are not removed and are simply distributed between taxa and if sufficiently repetitive will be set as saturated (see method section for details). Also, though CLARK-S and BBT both use multiple spaced seeds, our miBF did not use the same set of seeds as CLARK-S because of our restrictions on seed designs (see methods section). In addition, because our seed design does not affect memory usage of the miBF the same way it does in CLARK-S, we were also free to use longer seeds (Supp. Note S1).

Using the same simulated metagenomic datasets in the CLARK-S (40) paper we tried to replicate the results found within in addition to comparing CLARK and CLARK-S with BBT. Unfortunately, because the bacterial and viral NCBI databases have changed since the original CLARK-S publication, we had to omit read simulated from genomes that no longer have a corresponding species taxa in the database from the comparison, due to CLARK only selecting “Complete Genome [s]” and changes to the NCBI database demoting some genomes to draft status.

Nevertheless, since we omit the same reads in all runs and use the same reference sequences in each index database, our results will yield a fair comparison. There are two sets of simulated reads were generated as outlined in the CLARK-S paper; the difference between the “Default” and “Unambiguous” sets is the Unambiguous set does not have reads with all 32-mers shared between any 2 taxa IDs, but we note that, because reference database used has changed, this distinction may no longer hold completely true.

The default output of CLARK only produces a single best match, but CLARK-S can produce a secondary hit. Like CLARK-S, BBT can produce secondary hits in addition to a best hit, with the difference being that BBT can produce more than one secondary hits. The multi-matches parameterized to be filtered more stringently at the risk of losing the valid hits of a match. To this end, we compared the performance of each tool when it came to only the best hit in addition to comparing the results when multiple hits were considered.

In addition to default parameterizations of BBT (BBT Default), we tested it with parameterizations to increase specificity (BBT Specific) (For details on parameterization see Supp. Note S1). As expected, CLARK-S has higher sensitivity than CLARK in almost all cases, reproducing the results found in the original CLARK-S paper. The general trend shows that BBT Default has the highest sensitivity at the cost of precision, though this vast increase in sensitivity yields allows it to also yield the highest F1 score in all but one case (where BBT Specific has the highest F1). If considering the best hits, it is a toss-up between BBT Specific and CLARK in terms of highest precision, with BBT Specific generally yielding slightly higher sensitivity. However, when considering multi-hits, BBT Specific performs much better than all methods in terms of precision in all cases and shows comparable sensitivity to CLARK-S. For BBT Default and BBT Specific, only 7.5%, 10.14% have multi-hits respectively, and of that, a majority (60% and 74% respectively) of them only 2 possible hits (Supp. Fig S1).

Also included as part of the CLARK-S dataset was 3 negative control datasets totalling in 3,000,000 100 bp reads. Unexpectedly, some of the reads in the negative control datasets had mapped reads in both CLARK-S (6 reads) and Default BBT (3 reads). This is in contrast to the original CLARK-S paper where no reads were mapped in any of the negative controls. In addition, in BBT, we had expected few or no false positives because FPR was specified to be less than 10^−10^ (default parameters). Thus, we hypothesize this may be due to the difference in the databases, adding some new reference genomes that happen to have minor sequence similarity to those found in the negative control. Overall we feel this is not a cause for concern as this represents a very small minority of negative control reads. Finally, both CLARK and BBT Specific did not classify any negative reads.

On the same set of reads used in these experiments, we tested the runtime of each method at a differing number of threads (Fig. 7). We show that CLARK is the fastest tool when using a single thread, followed by BBT Default, BBT specific and finally CLARK-S. This trend generally follows when more threads are used, with the exception that BBT seems to scale better than CLARK, and will outperform CLARK when a least 8 threads within our tests. With the exception of the slow performance of CLARK-S, these methods remain within an order of magnitude of each other in performance if the same number of threads are used. In terms of memory, CLARK used 87GB, BBT used 91GB, and CLARK-S used 175GB. Database loading speed was not considered in this comparison but was dependent mostly on I/O and was quite comparable between the methods.

**Fig 6.**
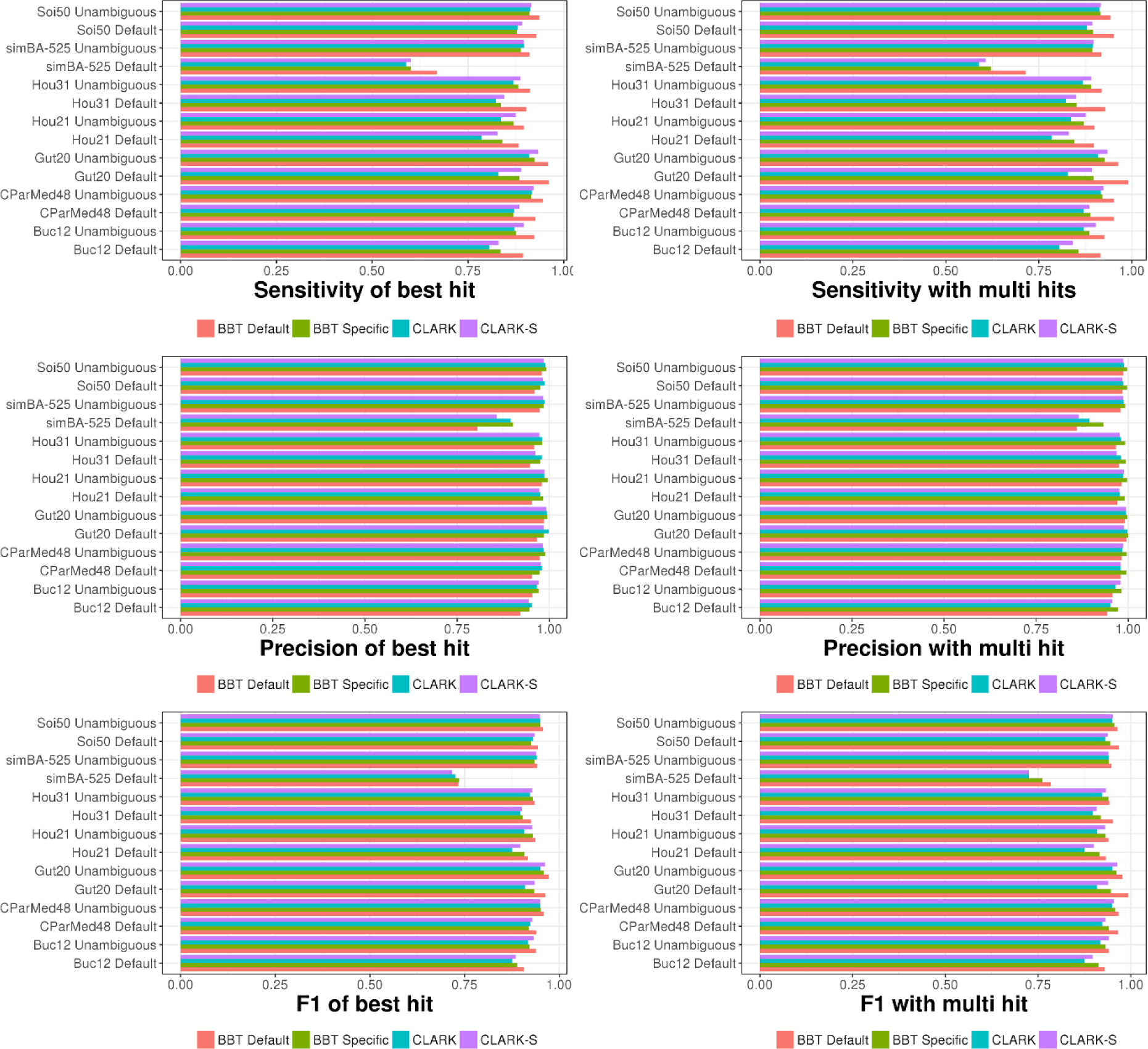
Precision and sensitivity comparison of CLARK, CLARK-S, BBT and BBT with parameterizations to increase specificity. On the right, only the best hit of a classification is considered correct, even if multiple species classifications are returned. On the left, reads are consider corrected if it multimaps to a correct species, even if multiple species are reported for the same read.

**Figure 7.**
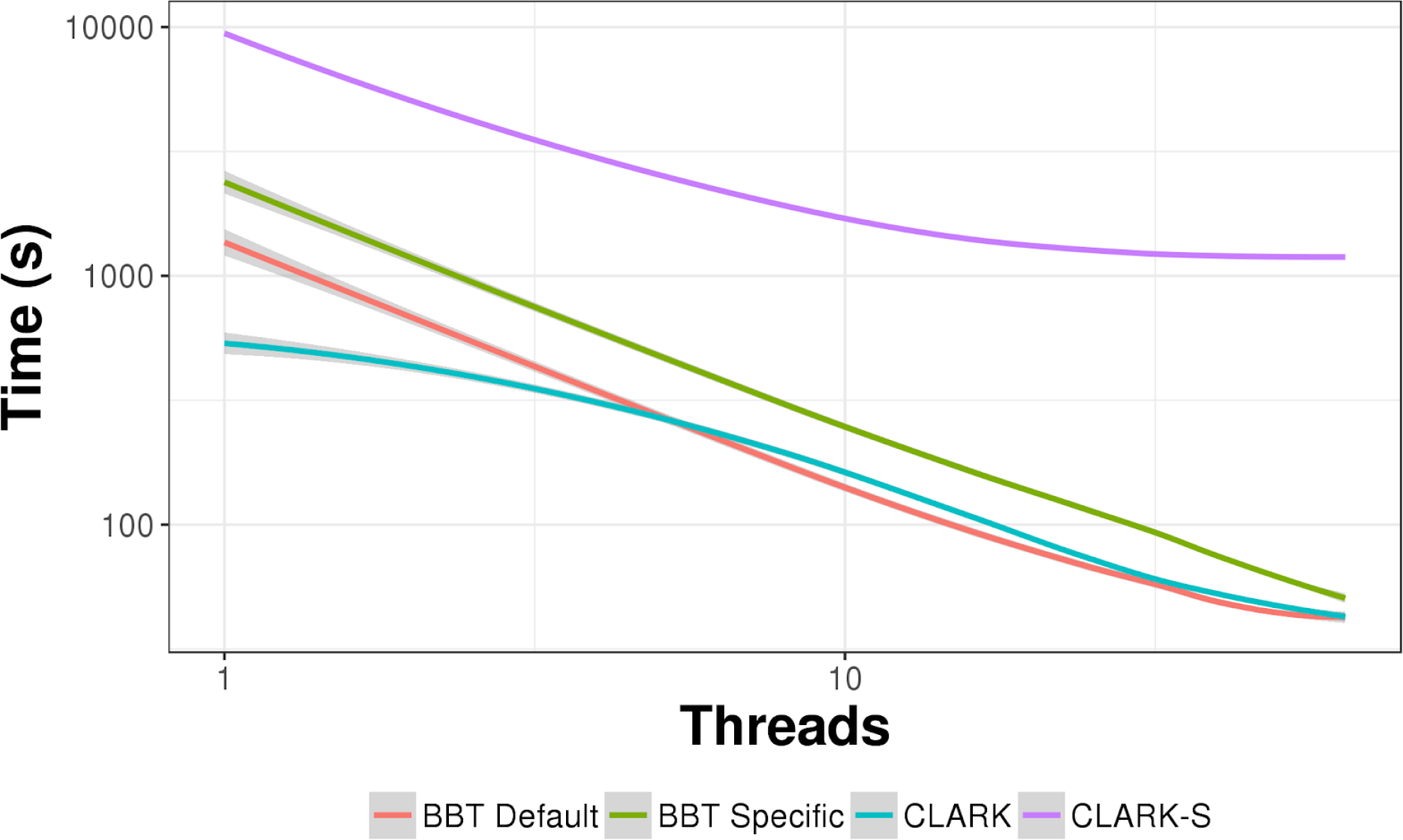
A runtime comparison of CLARK, CLARK-S, BBT and BBT with parameterizations to increase specificity at 1 to 64 threads. The axes are log scale, and under the situation of perfect scaling, the trend should follow a linear slope. Only classification time was considered, which does not include time taken to load the database from disk.

## DISCUSSION

We have presented the miBF, a probabilistic sequence classification data structure that can classify to multiple value-types and synergizes well with multiple spaced seeds. Query operations achieve a time complexity on par with a hash table based algorithms, but because it implicitly represents data, potentially uses less memory than hash table based algorithms (depending on the seed being hashed). Furthermore, seed size and complexity have no impact on memory usage on this data structure, expanding the potential for specialized, highly sensitive and specific spaced seed designs. In addition, we have formulated a model for calculating the FPR of classification using a probabilistic data structure given a sequence (rather than a single element query), which also leverages the frequency of which an ID is retrieved. Interestingly, this means miBFs with more IDs will result in lower FPRs. We showcase this highly sensitive generic classification tool implementation in two use cases.

The first use case was the recruitment of reads for targeted assembly. We show superior performance relative to BWA MEM in sensitivity and FPR at an order of magnitude faster time. The overall specificity of the matches was higher than BWA MEM but we note that in one target gene BWA MEM was better at disambiguating homologous sequences. However, for the purposes of targeted assembly, it is generally preferable to assign reads to both targets since missing sequences are more difficult to deal with than a few extra sequences. Furthermore, though the specificity of BWA MEM may be improved by including more of sequences to align to, to prevent off-target homologous alignment in cases where these sequences do not exist (e.g. incomplete or missing reference sequences), our results suggest that a spaced seed approach may be superior in terms of overall specificity.

The second use case was the classification of metagenomic sequences to the species level. Under default parameterization, we showed higher sensitivity than both CLARK-S and CLARK in classification when not only considering the multi-mapping reads but when considering just the best hits. To illustrate that the gains to sensitivity were not necessarily at the cost of precision, we also ran BBT with parameters to allow for more specific classifications. Under these parameters, we showed comparable specificity to CLARK and better specificity than CLARK-S for when only considering best hits, and higher specificity and sensitivity when considering multi-mapped reads. Our sensitivity gains were likely due to the use of a slightly sparser set of seeds with more seeds total (four vs three), and the because we do not filter out seed sequences that are shared between species. Despite the slight increase in sparseness, the specificity of the classification was maintained by using longer seeds (42bp, longer than what CLARK-S can currently use).

The runtime of BBT with default setting was comparable to CLARK, despite using multiple spaced seeds, and generally scales better than CLARK when more threads are used. When parameterized with more specific settings, BBT still remains less than twice the runtime of CLARK. Due to the implicit representation of the spaced seeds, BBT used only around half the memory CLARK-S and a similar amount of memory to CLARK despite using 4 different spaced seeds. The runtime of CLARK-S was expected to be slower than CLARK (10) but was much slower than expected as more threads were used. The runtime of CLARK-S more than an order magnitude slower than CLARK and BBT at a higher number of threads, suggesting that computation using multiple spaced seeds can be quite expensive if not carefully optimized.

### Future work

We use ntHash (26), a type of rolling hash (43) shown to be effective at computing hash values on nucleic acid sequences. ntHash is not yet been completely optimized for spaced seeds and hashing currently comprises of roughly 40-50% of our runtime. Currently, our implementation simply masks out positions where spaced seed no match positions exist, not effectively exploiting the properties of a rolling hash function. To improve performance it should be possible to improve ntHash to effectively use spaced seeds by adapting concepts from Girotto *et al* (44). Another improvement in terms of memory usage could be tuning the Bloom filter size by estimating the total number of distinct items for each spaced seed. This can be performed using ntCard (45), a streaming algorithm to estimate the *k*-mer abundance profile of large-scale genomics dataset, but with some modifications to adapt with spaced seeds rather than regular *k*-mers.

There remains a huge opportunity for further research into seed design with our data structure. Our data structure allows for unrestricted multiple spaced seed design relative by size or weight of the seed used. We investigated many seed designs and found although seed design does matter (Supp. Fig S3) according to metrics like overlap complexity (46), yet we still have not yet scratched the surface in terms of optimal spaced seed design. Optimal seed sensitivity computation NP-hard (47), and although faster approximations exist (48), they are still quite slow and infeasible because our seed can be any length. Optimal design for sequence classification is a function of sequencing error rate, homology detection tolerance, sequence length, and mutation/error types. Finally, if optimized seed hashing is implemented, it has been shown that some multiple spaced seed designs are more computationally efficient than others which is also a consideration to be made in seed design (44). In the end, we found that even randomly generated spaced seeds tend to perform well compared to *k*-mers (Supp. Fig S3) suggesting one can expect gains on sensitivity relative to *k*-mers without precise multiple spaced seed design. Thus, the seed used throughout the paper were randomly generated with a script provided as part of BBT after picking a weight and seed length that seems to work well for the use case. As seed design improves we expect the performance of our tool to improve.

### Conclusion

The formulation featured here for FPR reduction/calculation should be widely applicable to any probabilistic data structures when classifying sequencing data. Despite the complexity of using a tool based on a probabilistic data structure and spaced seeds, we expect this data structure and tool to be a valuable addition existing classification tools due to the impressive computational performance of the miBF as well as gains to classification sensitivity at a scalable memory usage. We hope that our work will invigorate research into spaced seed design because the length and weight of the seeds no longer have an impact on memory usage.

## DATA AVAILABILITY

A classification tool using our data structure is available as part of the BioBloom Tools GitHub repository (https://github.com/bcgsc/biobloom). Data structure implementation and library is available as part of our Bioinformatics Bloom Filter GitHub repository (https://github.com/bcgsc/btl_bloomfilter).

## ACKNOWLEDGEMENT

The authors would like to thank Rachid Ounit for providing support in the proper use of CLARK and CLARK-S, as well as providing insight into understanding the results produced using these tools using the datasets from the CLARK-S publication.

